# Synthetic Diffusion Tensor Imaging Maps Generated by 2D and 3D Probabilistic Diffusion Models: Evaluation and Applications

**DOI:** 10.1101/2025.02.21.639511

**Authors:** Tamoghna Chattopadhyay, Chirag Jagad, Rudransh Kush, Vraj Dharmesh Desai, Sophia I. Thomopoulos, Julio E. Villalón-Reina, Jose Luis Ambite, Greg Ver Steeg, Paul M. Thompson

## Abstract

Diffusion tensor imaging (DTI) is a key neuroimaging modality for assessing brain tissue microstructure, yet high-quality acquisitions are costly, time-intensive, and prone to artifacts. To address data scarcity and privacy concerns – and to augment the available data for training deep learning methods – synthetic DTI generation has gained interest. Specifically, denoising diffusion probabilistic models (DDPMs) have emerged as a promising approach due to their superior fidelity, diversity, controllability, and stability compared to generative adversarial networks (GANs) and variational autoencoders (VAEs). In this work, we evaluate the quality, fidelity and added value for downstream applications of synthetic DTI mean diffusivity (MD) maps generated by 2D slice-wise and 3D volume-wise DDPMs. We evaluate their computational efficiency and utility for data augmentation in two downstream tasks: sex classification and dementia classification using 2D and 3D convolutional neural networks (CNNs). Our findings show that 3D synthesis outperforms 2D slice-wise generation in downstream tasks. We present a benchmark analysis of synthetic diffusion-weighted imaging approaches, highlighting key trade-offs in image quality, diversity, efficiency, and downstream performance.

## I. Introduction

Diffusion tensor imaging (DTI [1]) is a powerful neuroimaging modality for studying the brain microstructure and anatomical connectivity. However, acquiring high-quality DTI scans is costly, time-consuming, and susceptible to artifacts. The growing demand for large-scale, high-quality neuroimaging datasets has driven interest in synthetic data generation to supplement real acquisitions. Synthetic DTI generation can help mitigate data scarcity, and enhance the accuracy and generalization ability of models trained on the real and synthetic data. With appropriate safeguards, synthetic data may also offer a way to overcome data privacy issues in medical research by generating realistic brain scans that do not belong to any specific individual, yet represent the true variation in the population. Recent advances in deep generative models, particularly denoising diffusion probabilistic models (DDPMs) [2,3,4], have yielded state-of-the-art results in synthesizing high-fidelity medical images. Compared to generative adversarial networks (GANs) [5] and variational autoencoders (VAEs) [6], DDPMs typically offer better mode coverage, improved sample diversity, and more stable training dynamics, making them well-suited for generating complex medical imaging data. GAN-based methods often suffer from mode collapse, limiting their ability to represent the full variability of brain structures. VAEs, on the other hand, struggle with blurriness due to the constraints imposed by their latent space. DDPMs [7,8,9,10] use a sequential denoising process that produces high-resolution, diverse, and realistic synthetic brain scans, making them ideal for applications in neuroscience. When used for medical image synthesis [11,12,13,14], DDPMs can preserve fine-scale anatomical features and maintain statistical properties of real medical scans. Even so, 3D volumetric images can be more challenging to generate than 2D slices given the complex spatial relationships between anatomical structures and the need to maintain neuroanatomical consistency across slices. In the current work we focus on DTI-derived maps of mean diffusivity (MD), a widely-used metric that is sensitive to brain aging [15], Alzheimer’s disease [16] and Parkinson’s disease [17]. This metric is an important marker of brain integrity (which tends to decline with old age and disease) and is a source of image contrast not evident on standard anatomical MRI.

Recent research has examined both 2D and 3D approaches to medical image synthesis, leading to questions about which approach is optimal when considering downstream tasks. Many synthetic neuroimaging studies focus on 2D slice-based approaches as they are computationally more efficient and large-scale 2D datasets are available. However, 3D generation preserves spatial coherence across slices, which may be crucial for downstream tasks such as disease classification, shape modeling, and volume quantification. In this work, we train 2D and 3D diffusion models to generate synthetic diffusion tensor imaging (DTI) mean diffusivity (MD) maps—a widely used scalar metric for assessing the brain’s microstructure in clinical and research settings. We evaluate training strategies under constraints of limited data, computational resources, and memory efficiency. This includes optimizing model architecture, training duration, and incorporating a latent diffusion framework to enhance performance. We also evaluate the value of using synthetic data to augment the training data for downstream classification tasks (sex classification and dementia detection) using 2D CNNs vs. 3D CNNs. We investigate whether generating a single slice is sufficient for learning discriminative features, or if full 3D brain synthesis is needed to provide information that is needed for the task. Our findings provide a benchmark for the practical application of synthetic diffusion-weighted images in radiology and neuroscience.

Our contributions are threefold: (1) we compare 2D and 3D DDPM architectures that are optimized for DTI-MD synthesis, (2) we analyze the quality, fidelity, and diversity of data generated from slice-wise and volume-wise generative models, and (3) we evaluate the effectiveness and computational efficiency of each approach for supporting downstream classification tasks, offering insights into the practical utility of different generation strategies.

## II. Data and Preprocessing

Diffusion tensor imaging (DTI) is widely used to evaluate brain tissue microstructure *in vivo*. As an extension of conventional MRI, it uses ‘diffusion weighting’ [18] to enhance sensitivity to water diffusion rates in multiple directions [1]. The diffusion tensor model models the local diffusion of water in brain tissue using a spatially varying tensor (**Fig. 1**). This diffusion process can also be represented as a 3D Gaussian distribution at each voxel - diagonal elements of the DT represent the variance (or the squared magnitude of diffusion) along the principal axes of diffusion; off-diagonal elements represent the covariance between displacements in different directions. Although more complex diffusion models have also been used in research (such as DKI[19], NODDI[20] and MAP-MRI[21]), key quantitative metrics derived from DTI include fractional anisotropy (FA) and three diffusivity measures—mean (MD), axial (AxD), and radial (RD) diffusivity. These metrics capture the tensor’s geometric properties, reflecting water diffusion along its principal axes. Notably, MD can detect variations in cellular density, edema, and necrotic processes, making it a critical marker in neurodegenerative conditions such as Alzheimer’s [22] and Parkinson’s [17] diseases. The standard formula for MD, in terms of the diffusion tensor eigenvalues, is:

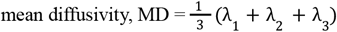

**Fig. 1.**
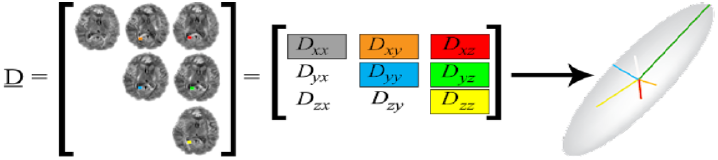
Diffusion Tensor Components. In the simplest case, the diffusion tensor represents a 3D Gaussian model of water diffusion, modeled using a diffusion ellipsoid, or a 3×3 matrix representing diffusion rates in different directions; this tensor can be rotated to assume a diagonal matrix form with only 3 diagonal components - the eigenvalues - which represent the relative diffusion tendencies in each of 3 principal directions. The mean diffusivity is approximately a spherical average of the diffusion magnitudes in all directions.

In our study, we analyzed the Cambridge Centre for Aging and Neuroscience (Cam-CAN) dataset [23], chosen for its broad age distribution (range: 18-88 years). We analyzed diffusion MRI data from 652 cognitively normal participants (mean age: 54.3 ± 18.6 years; 322 females, 330 males). The diffusion MRI preprocessing steps are detailed in [24,25]. The 3D generative models in this study were trained using 3D DTI-MD maps from Cam-CAN, whereas the 2D models were trained using the middle axial slice from the dataset. For 3D, the final image sizes for the model were 64×64×64, whereas, for 2D, the final image slice was 128×128. For downstream tasks, we included data from the Alzheimer’s Disease Neuroimaging Initiative [26]. We used diffusion MRI data from 332 subjects (mean age: 74.61 ±8.14 (SD) years; 163 F/169 M). For this dataset, the diagnostic distribution was 199 healthy controls and 133 participants with dementia.

### II. Deep Learning Architectures

In this paper, we distinguish *diffusion MRI*–a medical imaging technique to measure water diffusion in biological tissue–from *probabilistic diffusion models*, a type of generative model used in deep learning, which is based on statistical thermodynamics and Langevin diffusion. Langevin dynamics [27] simulates a random walk that gradually transitions from a noise distribution to a data distribution, typically controlled by a differential equation that incrementally removes noise. Diffusion models approximate the underlying data distribution by iteratively refining a learned distribution using a neural network. During training, noise–sampled from a normal distribution– is progressively added to an image following a predefined noise schedule, such as a linear scheduler, where noise is introduced in discrete steps. Architectures such as the diffusion pixel-space-based DDPM [28] and latent diffusion models (LDM [2,29]) commonly use an enhanced U-Net [30] structure with self-attention mechanisms. Additionally, discrete time steps can include conditioning inputs (such as a text prompt that specifies the type of image to generate) via cross-attention mechanisms embedded within ResNet blocks. The primary goal of a diffusion model is to estimate the noise added at a random step, minimizing a regression loss—typically mean squared error (MSE). Once trained, the model generates synthetic scans using different sampling strategies, such as DDPM or its deterministic counterpart, the denoising diffusion implicit model (DDIM [31]), with a predefined number of steps. Through this iterative process, the model transforms random noise into a realistic image that closely aligns with the learned distribution. The choice of sampling scheduler impacts both the quality of the images produced and the computational efficiency. Unlike DDPMs, LDMs (**Fig. 2**) typically use autoencoders to project images into a lower-dimensional latent space before applying diffusion processes, reducing the computational burden of training deep models on high-dimensional 3D MRI scans. The autoencoder in LDMs is trained using a combination of multiple loss functions, including, reconstruction loss, perceptual loss, KL divergence loss, and adversarial loss. In this study, we trained both LDM and DDPM architectures for comparison. A key consideration in cr eating synthetic medical images is the choice of 2D slice-wise versus 3D volume-wise diffusion models. Both architectures share fundamental components, but their implementations and computational trade-offs differ considerably.

**Fig. 2.**
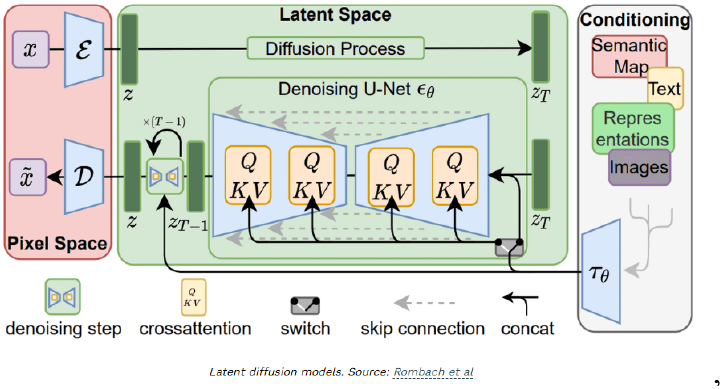
Latent Diffusion Model Architecture, reproduced from Rombach et al. [2].

#### 2D Diffusion Models

These models generate individual slices of a brain scan independently. They are computationally efficient and require fewer resources. They can be trained on datasets that lack full 3D volumes, for applications where individual slices are sufficient. However, they do not enforce anatomical consistency across generated slices, which may limit their utility for tasks requiring quantification of structures in 3D.

#### 3D Diffusion Models

Instead of generating individual slices, 3D models operate on entire brain volumes, generating data with spatial consistency across slices. These models require more memory and computational resources, as they need to process batches of 3D medical images. While these models capture 3D spatial context, training and inference require more time and memory.

Following standard practices for evaluating generative models, we quantitatively assessed the realism and diversity of the generated synthetic images using established performance metrics. The evaluation was conducted over 500 image pairs, using maximum mean discrepancy (MMD) [32] and multi-scale structural similarity index (MS-SSIM) to quantify similarity. A lower MMD score indicates that the synthetic data distribution closely aligns with the real data distribution, while a higher MS-SSIM value reflects greater structural similarity. To assess realism, we compared real and synthetic images within the same class using MMD and MS-SSIM. For diversity, MS-SSIM was used to measure differences between two distinct synthetic images generated under the same class conditions, with real image pairs serving as a reference for comparison.

The 3D CNN architecture (**Fig. 3**) consisted of four convolutional layers with a 3×3 kernel size, followed by a 1×1 convolutional layer and a final dense layer. Each layer used the ReLU activation function and Instance Normalization to stabilize training. To mitigate overfitting, dropout layers with a dropout rate of 0.5 were incorporated after the 2nd, 3rd, and 4th convolutional layers, alongside a 3D average pooling layer with a 2×2 kernel. The model was trained using the Adam optimizer [33] with a learning rate of 10^−4^, which decayed exponentially at a rate of 0.96. Training was conducted for 100 epochs with a batch size of 8, using mean squared error as the loss function. Early stopping and dropout regularization were employed to reduce overfitting. Model performance was first evaluated on a sex classification task–a common benchmarking task as ground truth is known–using balanced accuracy and F1 score. For the 2D CNN version, all layers were adapted to operate in 2D by replacing 3D convolutional and pooling operations with their 2D counterparts, while maintaining the overall architectural structure. After registering all images to a common template, the dataset was divided into training, validation, and testing subsets in an approximate 70:20:10 ratio. Two experimental setups were used: (1) starting with a fully real-image training set and progressively replacing real images with synthetic ones, and (2) maintaining a constant number of real training images while adding synthetic samples. These experiments were conducted for both DDPM- and LDM-generated images to assess their impact on downstream classification performance.

**Fig. 3.**
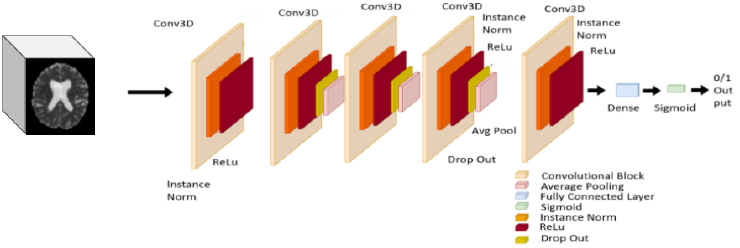
3D CNN Architecture that we trained for downstream tasks.

**Fig. 4.**
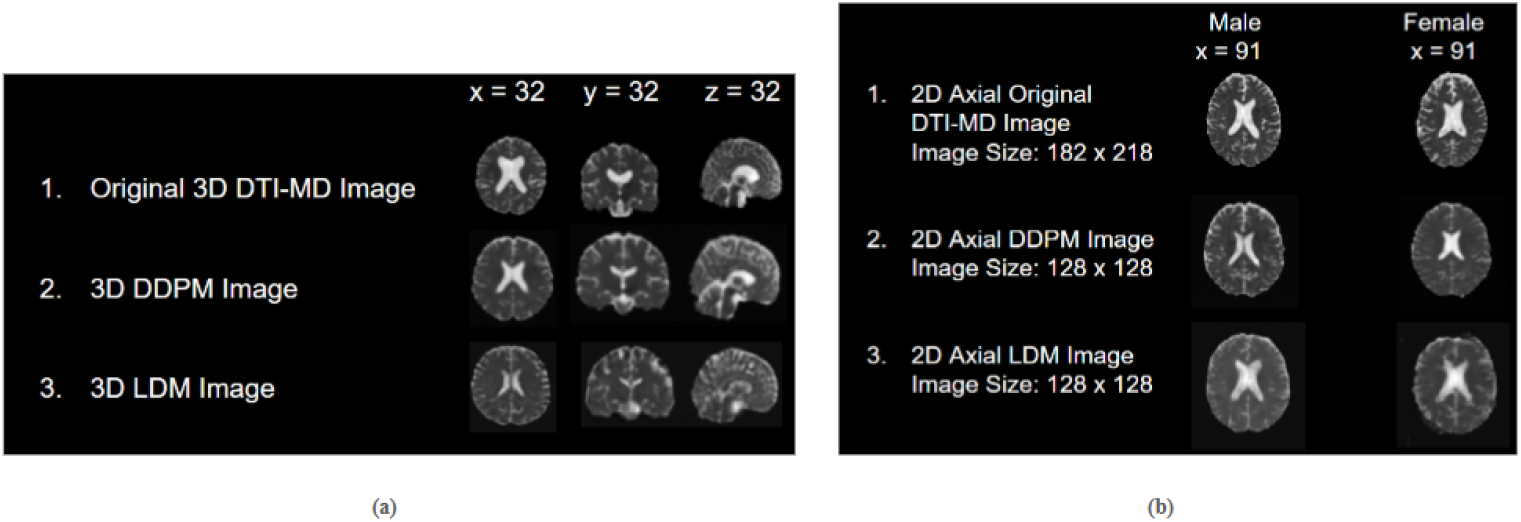
**(a)** The first row shows the original real DTI-MD maps from a randomly selected participant in the Cam-CAN dataset. The next two rows show synthetic DTI-MD maps generated by the DDPM and LDM. Notably, some sulcal features (fissures) in the cortex are visible, even though the input data is only sampled with 2-mm isotropic voxels (a commonly-used spatial resolution for DTI). The slices in figure are all for male participants. **(b)** The first row shows the middle axial slice used for 2D versions of the model. The next two rows show the middle axial slice synthetically generated by DDPM and LDM, respectively.

**Fig. 5.**
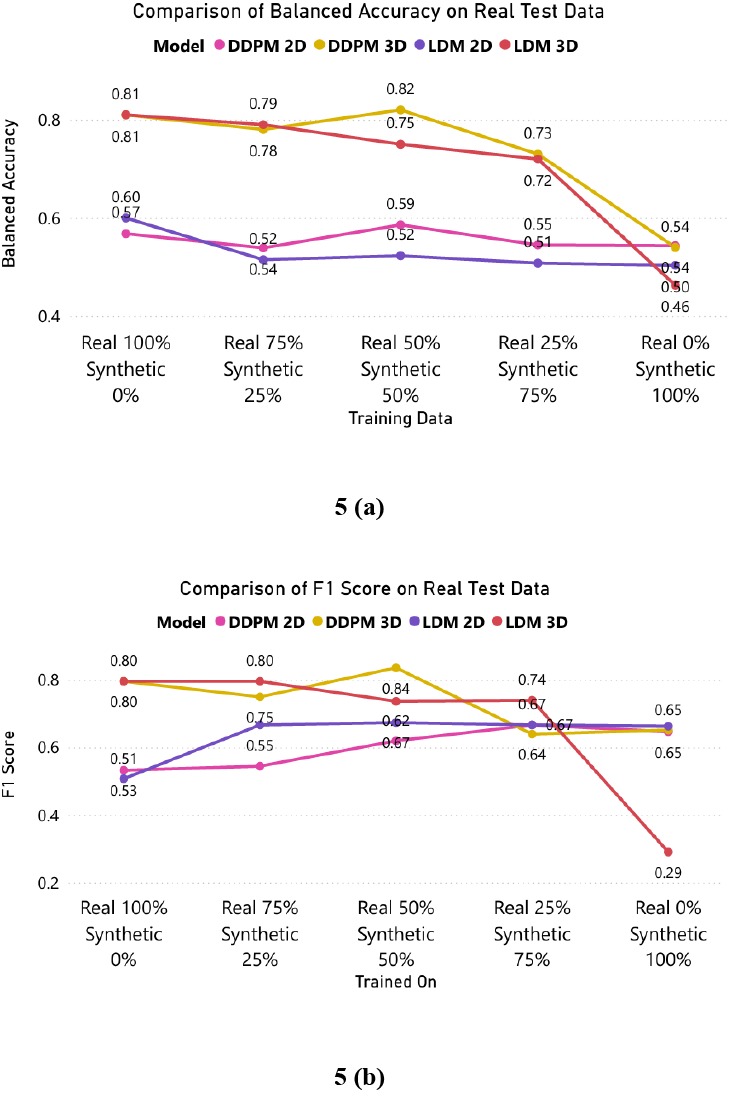
**(a)** Comparison of Balanced Accuracy on Real Test Data. **(b)** Comparison of F1 Score on Real Test Data. Particularly for the 3D models, the results are generally in line with the expectation that performance will decline as less real data is used, and more synthetic data is used. In the limit where only synthetic data is used for training, performance declines for the 3D models.

## IV. Experimental Results and Discussion

Evaluations of the synthetic DTI-MD maps generated by the different models, compared to real scans, are summarized in **Table 1**. Best performances are highlighted in **bold**. For both male and female participants, LDM models used to generate synthetic images consistently outperformed DDPMs on realism metrics, particularly in terms of MMD and MS-SSIM (Real vs. Synthetic). The best realism score (lowest MMD) was achieved by the 3D LDM model for both male (0.0130±0.009) and female (0.0086±0.006) participants, suggesting the value of these models in preserving anatomical consistency. Similarly, 2D LDM models achieved the highest MS-SSIM (Real vs. Synthetic) values, further supporting their advantage in generating images that closely resemble real data. However, when evaluating the diversity of generated samples, DDPM models performed best, particularly in the MS-SSIM (Synthetic vs. Synthetic) metric, which measures intra-model variability. The 2D DDPM achieved the highest diversity for male subjects (0.7351±0.201), while the DDPM 3D model showed strong diversity for female participants (0.781±0.175). These findings suggest that DDPMs may produce a broader range of synthetic samples, perhaps due to their direct pixel-space generation, which avoids the compression-induced homogenization observed in latent diffusion models.

**Table 1.** Realism and Diversity Test Metrics for the DDPMs and the LDMs. The best model performance for each metric is shown in **bold**. The MMD is computed from pairs drawn from the real and synthetic data distributions; lower values are better. SSIM is also a measure of realism, where higher values are better when real and synthetic images pairs are compared. SSIM can be adapted to measure diversity in synthetic images by interpreting low similarity as high diversity. A high SSIM (close to 1) indicates that two images are very similar, whereas a lower SSIM indicates greater dissimilarity. To measure diversity with SSIM, lower SSIM scores would suggest greater diversity between pairs of images. In the last two columns, the diversity of the real data is considered as a reasonable lower bound for this metric.

| Model | Image Size | Train Epochs | Scheduler | Realism: MMD (Real vs Synthetic)↓ | Realism: MS-SSIM (Real vs Synthetic)↑ | Diversity: MS-SSIM (Synthetic vs Synthetic)↓ | Diversity: MS-SSIM (Real vs Real)↓ |
| --- | --- | --- | --- | --- | --- | --- | --- |
| **Cam-CAN Male Subjects** |  |  |  |  |  |  |  |
| **DDPM 2D** | 128x128 | 1000 | DDPM | 0.0235 ± 0.015 | 0.7193 ± 0.086 | 0.7834 ± 0.087 | 0.7153 ± 0.054 |
| **DDPM 3D** | 64x64x64 | 2000 | DDPM | 0.0283 ± 0.029 | 0.5949 ± 0.245 | **0.7351 ± 0.201** | 0.6522 ± 0.237 |
| **LDM 2D** | 128x128 | 80 (AE) + 1000 | DDPM | 0.0175 ± 0.008 | **0.7395 ± 0.054** | 0.8052 ± 0.052 | 0.7153 ± 0.054 |
| **LDM 3D** | 64x64x64 | 80 (AE) + 2000 | DDPM | **0.0130 ± 0.009** | 0.6428 ± 0.216 | 0.7813 ± 0.157 | 0.6522 ± 0.237 |
| **Cam-CAN Female Subjects** |  |  |  |  |  |  |  |
| **DDPM 2D** | 128x128 | 1000 | DDPM | 0.0170 ± 0.009 | **0.7551 ± 0.062** | 0.8337 ± 0.054 | 0.7285 ± 0.049 |
| **DDPM 3D** | 64x64x64 | 3000 | DDPM | 0.0350 ± 0.035 | 0.6530 ± 0.214 | 0.7805 ± 0.175 | 0.6974 ± 0.188 |
| **LDM 2D** | 128x128 | 80 (AE) + 1000 | DDPM | 0.0241 ± 0.007 | 0.6462 ± 0.039 | **0.7482 ± 0.075** | 0.7285 ± 0.049 |
| **LDM 3D** | 64x64x64 | 80 (AE) + 2000 | DDPM | **0.0086 ± 0.006** | 0.7441 ± 0.115 | 0.8886 ± 0.064 | 0.6974 ± 0.188 |

The overall diversity scores are relatively high, which aligns with the expectation set by the diversity scores for real images, providing a reasonable lower bound for this metric. LDMs may score higher for realism as they can model fine-grained spatial structures, perhaps due to their latent-space representation and perceptual loss functions. Conversely, the greater diversity observed in DDPM-generated samples may result from their less constrained generative process, which may capture a wider range of anatomical variations. This trade-off between realism and diversity highlights the importance of selecting the appropriate generative framework based on the specific requirements of downstream tasks.

For the first downstream experiment, we assessed the performance of CNN models trained on a combination of real and synthetic DTI-MD maps for the task of sex classification. We evaluated balanced accuracy and F1 scores on real test dataset across the four generative models: 2D and 3D DDPM, and 2D and 3D LDM. The experiment followed a progressive data replacement strategy, where the proportion of synthetic data in the training set was systematically reduced in increments of 25%, with real data from the Cam-CAN dataset replacing it, in each successive training run. From our experiments, we see that on real test data, the 3D synthetic models outperformed the 2D models. The 3D DDPM performed best, achieving a peak balanced accuracy of 0.82 when trained with 50% real and 50% synthetic training data. This suggests that incorporating a small fraction of synthetic data does not degrade performance and may help the model to generalize. The 2D models had accuracies ranging from chance level to 0.6. 3D generative models outperformed their 2D counterparts, suggesting the importance of preserving spatial dependencies in DTI-MD maps for the downstream task of sex classification. The 3D DDPM model, in particular, maintains strong performance across different training data compositions. In contrast, the 2D models struggle, with their accuracy near chance levels, suggesting that they fail to retain the spatial context needed for reliable classification. This substantial performance gap suggests that 3D volumetric coverage is crucial, at least for this classification task.

In the second downstream experiment, we assessed the performance of CNN models on the sex classification task while keeping the amount of real training data constant but now *increasing* the proportion of synthetic data in increments of 25% across successive runs. We evaluated balanced accuracy and F1 scores on real test data for four models: 2D and 3D DDPM, and 2D and 3D LDM. The 3D synthetic models outperformed the 2D models. In **Figures 6(a,b)**, we did not find evidence that the synthetic data helped or harmed performance, at least for this task. Even so, the 3D DDPM model stands out, maintaining strong performance across different training data compositions.

**Fig. 6.**
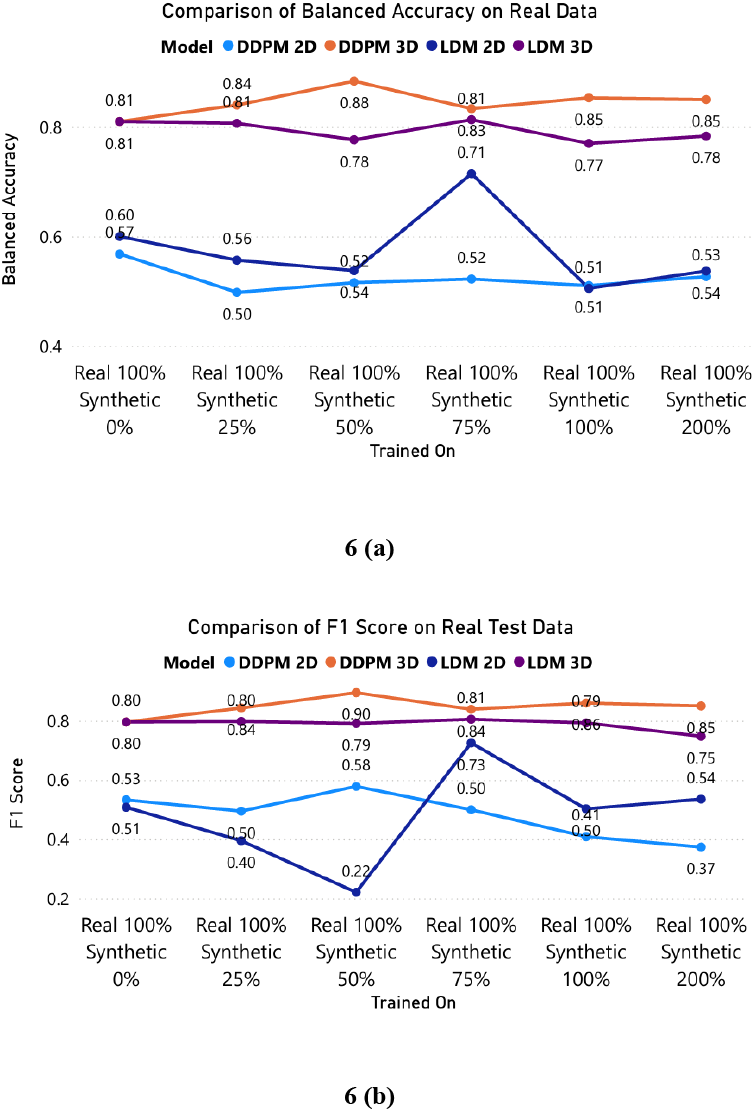
**(a)** Comparison of Balanced Accuracy on Real Test Data. **(b)** Comparison of F1 Score on Real Test Data.

For the third downstream task, we chose transfer learning for dementia classification. Sex classification has been used as an effective pretraining task for dementia classification as it can help models to learn generic neuroanatomical features from brain imaging data [34,35,36]. As sex differences in brain structure and in diffusion characteristics are well-documented, training a model to distinguish male and female brains helps it to capture global and regional features of brain morphology (or, in this case, of microstructure). These learned representations can then be transferred to dementia classification, where subtle structural changes, such as cortical atrophy and white matter degeneration, can be used to distinguish patients with dementia from healthy elderly controls. By leveraging the knowledge gained from sex classification, models can achieve better feature extraction and improved performance in detecting dementia-related abnormalities. The results in **Table 2** indicate the impact of pretraining on sex classification for the downstream task of dementia classification. The highest 3D balanced accuracy (0.808) is achieved when using an LDM model pretrained solely on 100% synthetic data, suggesting that synthetic data can be useful in enhancing performance. Comparatively, the highest 2D balanced accuracy (0.647) is also observed with LDM trained on synthetic data, further reinforcing this trend. DDPM-based models generally perform slightly worse than LDM: balanced accuracy values range between 0.727 and 0.797 for 3D, with lower values for 2D. Additionally, models trained with both real and synthetic data do not consistently outperform those trained on only one type. The results show that 3D outperforms 2D, but the best model only slightly improves balanced accuracy over no pretraining. This could be because although the pretraining model is trained on a task in the same domain, the datasets are different. Cam-CAN only has controls whereas ADNI has participants with dementia, so there is an inherent difference between the two datasets. Moreover, the amount of training data for downstream tasks is quite low as well as the amount of training data for the pretraining task. More training data may lead to better performance.

**Table 2.** Dementia Classification Results, using sex classification as pretraining task. The best model performance is shown in **bold**. 2D indicates that the training dataset for Dementia Classification consists of 2D middle axial slice, whereas 3D indicates that the whole volume was used for training. ✓indicates pretraining was done whereas x means that pretraining was not done. - indicates that no data from that particular subset of real or synthetic was used for training the sex classifier. The percentages indicate the amount of data used for pretraining - 200% indicates double the amount of synthetic data in training.

| PreTraining | Model | Trained on |  | 2D |  | 3D |  |
| --- | --- | --- | --- | --- | --- | --- | --- |
|  |  | Real Data | Synthetic Data | Balanced Accuracy | F1 Score | Balanced Accuracy | F1 Score |
| x | - | - | - | 0.607 | 0.467 | 0.798 | 0.732 |
| ✓ | DDPM | 100% | - | 0.588 | 0.435 | 0.727 | 0.656 |
| ✓ | DDPM | 100% | 100% | 0.595 | 0.471 | 0.760 | 0.690 |
| ✓ | DDPM | 100% | 200% | 0.546 | 0.401 | 0.797 | 0.736 |
| ✓ | DDPM | - | 100% | 0.585 | 0.461 | 0.741 | 0.678 |
| ✓ | LDM | 100% | - | 0.616 | 0.488 | 0.749 | 0.676 |
| ✓ | LDM | 100% | 100% | 0.636 | 0.519 | 0.798 | 0.732 |
| ✓ | LDM | 100% | 200% | 0.627 | 0.530 | 0.760 | 0.690 |
| ✓ | LDM | - | 100% | 0.647 | 0.568 | **0.808** | **0.746** |

## V. Conclusion and Future Work

In this work, we comprehensively evaluated 2D and 3D DDPMs and LDMs for generating synthetic DTI-MD scans. We evaluated how synthetic data from these models may affect downstream performance, specifically for the tasks of sex classification and dementia classification using both 2D and 3D convolutional neural networks (CNNs). The full 3D model performs better which is in line with the expectation that 3D data will improve performance.

This supports the hypothesis that generating the entire 3D brain volume provides richer, more contextual information for downstream classification tasks. To add context to the current work, there are ongoing related efforts using probabilistic diffusion models for image enhancement (super-resolution) and image translation (generation of new contrasts). Relatively few studies have focused on generation of diffusion MRI, with some exceptions [37,38,39,40,41]. Our findings highlight the promise of diffusion-based generative models in augmenting limited datasets and facilitating a range of analytic workflows. Future research will evaluate the effect of increasing the diversity of synthetic data, test the models on larger, more diverse, multi-site datasets, and examine additional downstream applications, such as prognosis prediction, to demonstrate their value in biomedical research.

## Acknowledgments

This work was supported by NIH NIA grants U01AG068057 (‘AI4AD’) and R01AG081571 (‘Federate AD’).

